# Systematic analysis of gut bacterial carcinogen metabolism and its functional consequences

**DOI:** 10.1101/2024.05.20.595058

**Authors:** Boyao Zhang, George-Eugen Maftei, Bartosz Bartmanski, Michael Zimmermann

## Abstract

Organic carcinogens, in particular DNA-reactive compounds, contribute to the irreversible initiation step of tumorigenesis through introduction of genomic instability. Although carcinogen bioactivation and detoxification by human enzymes has been extensively studied, carcinogen biotransformation by human-associated bacteria, the microbiota, has not yet been systematically investigated. We tested the biotransformation of 68 mutagenic carcinogens by 34 bacterial species representative for the upper and lower human gastrointestinal tract and found that the majority (41) of the tested carcinogens undergo bacterial biotransformation. To assess the functional consequences of microbial carcinogen metabolism, we developed a pipeline to couple gut bacterial carcinogen biotransformation assays with Ames mutagenicity testing and liver biotransformation experiments. This revealed a bidirectional crosstalk between gut microbiota and host carcinogen metabolism, which we validated in gnotobiotic mouse models. Overall, the systematic assessment of gut microbiota carcinogen biotransformation and its interplay with host metabolism highlights the gut microbiome as an important modulator of exposome-induced tumorigenesis.

## Introduction

Organic mutagens are a class of carcinogens that can cause DNA mutations and induce tumor development^1^. These compounds are found in various sources such as food (*e.g.*, heterocyclic amines from cooking processes^2^) and the environment (*e.g.*, nitrosamines from incineration processes, cigarette smoke, and chloramine water sanitation^3^). Following uptake of such organic mutagens by the human body, multiple metabolic enzyme systems (*e.g.*, cytochrome P450 and glutathione S-transferases) chemically alter these compounds in various biotransformation reactions, such as oxidation, hydrolysis, and conjugation. Such metabolism renders the chemicals more hydrophilic and hence easier to eliminate from the body, which in turn reduces their systemic exposure. Although carcinogen biotransformation ideally detoxifies compounds, certain mutagens are converted into more genotoxic forms by host metabolic reactions and hence, biotransformation increases their mutagenic activity (known as bioactivation) and the hazard to the host^4,5^.

In addition to human enzymes, the microbes residing in the human gastrointestinal tract (known as the ‘gut microbiota’) encode at least 100-fold more genes than the human genome, and increasing evidence suggests a role of microbiome-encoded enzymes in xenobiotic biotransformation^6–9^. For example, biotransformation of clinical drugs by human gut bacteria was found to be a common phenomenon^8^. On the other hand, numerous studies have associated cancer development with altered microbiome composition and specific taxonomic features^10,11^. However, mechanistic insights into how gut bacteria impact tumorigenesis remain mostly anecdotal. Intriguingly, some studies suggested that lower incidences of cancer development could be due to bioaccumulation and degradation of carcinogenic compounds by specific gut bacterial species^12–15^. However, there is a lack of systematic understanding of the metabolic potential of gut bacteria for various organic carcinogens and the functional consequences of microbiota mutagen metabolism for the host. Furthermore, given that both human and bacterial enzymes can either aid detoxification or bioactivation of carcinogens, it is necessary to assay the interactions between host and microbial metabolism of carcinogens to predict their fates and activities in the human body.

In this study, we screened a diverse range of environmental organic mutagens for their biotransformation by representative human gut bacteria in a high-throughput manner to assess the capacity of gut bacteria to metabolize organic mutagens. We then quantitatively assessed the changes in chemical mutagenicity of specific compounds upon bacterial metabolism through the coupling of bacterial biotransformation assays to Ames mutagenicity tests. Further, we combined human microsome and bacterial biotransformation assays of mutagens with mutagenicity tests to gain insights into the functional consequences of the host-microbial metabolic crosstalk on mutagenicity. Finally, we used gnotobiotic mouse models to demonstrate that the crosstalk between host and bacterial metabolism observed *in vitro*, also reflects metabolic interactions of carcinogens *in vivo*.

## Results

### Environmental organic mutagens are widely metabolized by gastrointestinal bacteria

To systematically assess the biotransformation of mutagenic carcinogens (referred to as ‘mutagens’) by human-associated bacteria, we assembled a panel of 68 diverse organic compounds with mutagenic activity. The chemicals were selected based on their mutagenicity in Ames tests with or without prior microsomal bioactivation documented in the EURL ECVAM Genotoxicity and Carcinogenicity Consolidated Database^16^ (Table S1). Cytochrome P450 enzymes captured in microsomal liver preparations are a major contributor to bioactivation of carcinogens and typically used to test for carcinogen bioactivation^4^. The selected chemicals covered a diverse range of origins, such as cigarette smoke, industrial processes and clinical drugs (Fig. 1A), and physicochemical properties, such as molecular weight and logP values (Fig. 1B-C).

**Figure 1.**
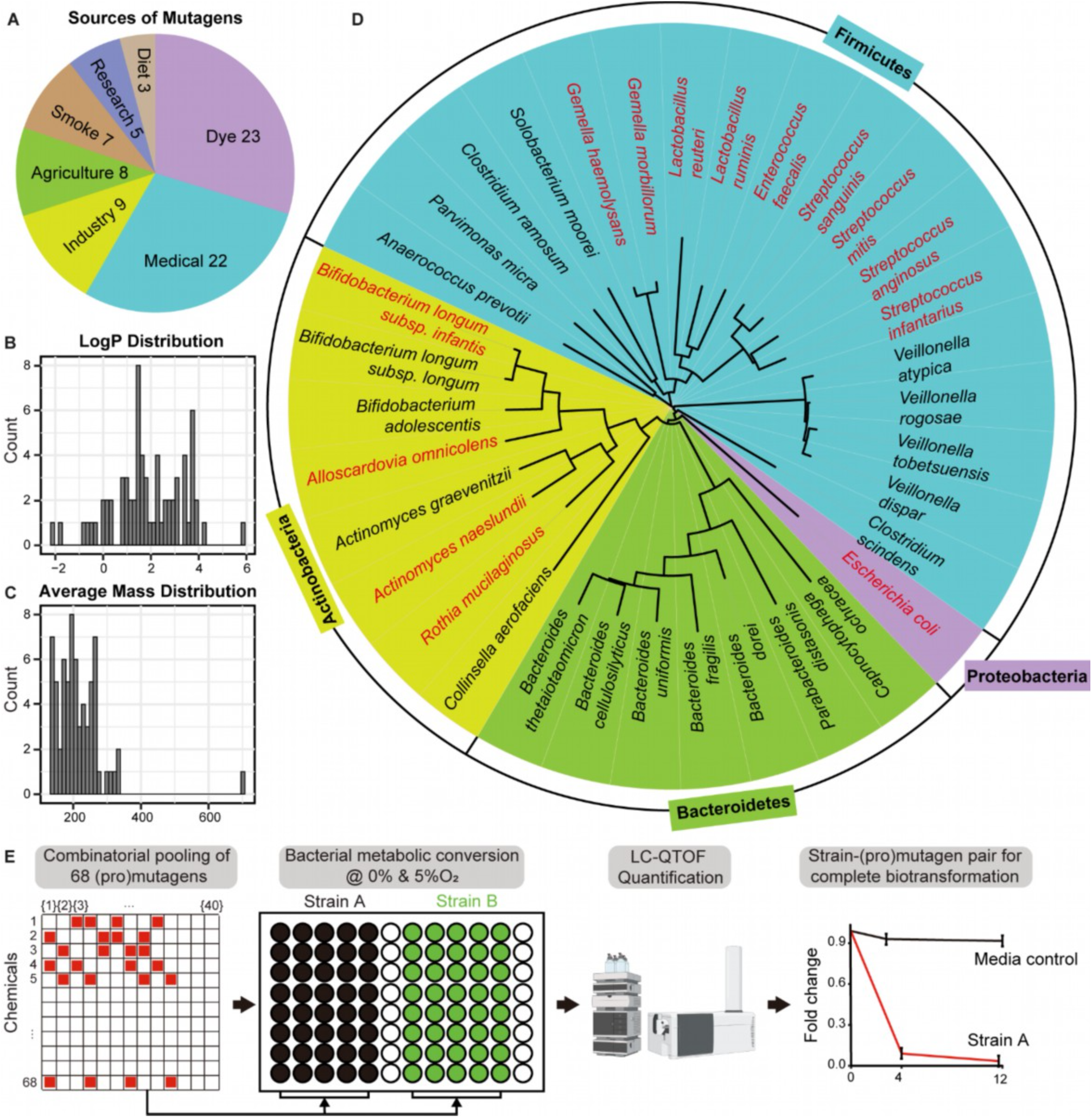
Carcinogen and bacterial species used in the biotransformation assay. **A.** Sources of mutagenic carcinogens tested in this study (some compounds belong to multiple categories). LogP (B) and monoisotopic mass (C) distribution of the 68 tested chemicals. **D.** Bacterial species tested represent different phyla commonly found in the human intestinal tract. Species solely tested under anaerobic condition are depicted in black and species tested under both anaerobic and microaerobic conditionsn are shown in red. **E.** Experimental pipeline to systematically test gut bacterial biotransformation of organic carcinogens.

To assay gut microbial biotransformation of the assembled mutagens, we selected 34 phylogenetically representative bacterial species from the human gastrointestinal tract (GIT), including bacteria isolated from the oral cavity, the small intestine and the large intestine (Fig. 1D and Table S2). To account for oxygen gradients that exist between the upper and lower intestine and between the gut lumen and the epithelium^17^, we assayed all 34 species under anaerobic conditions and the 14 nonobligate anaerobe species of our panel also under microaerobic (5% O_2_) culture conditions. Mutagens were assembled in 40 combinatorial pools, where each chemical was assayed in four independent replicates at 14.3 uM and each chemical was pooled with any of the other chemicals not more than twice. We then incubated each bacterial culture with each pool of chemicals and monitored carcinogen biotransformation by liquid chromatography coupled to mass spectrometry (LCMS) at 0, 4, and 12 hours of incubation (Fig. 1E). We found that a total of 41 of the 68 tested mutagens were biotransformed (>75%, adj p-value: 0.05) after 12 hours by at least one bacterial species (Fig. 2A-B and Table S3). We also found marked inter-species differences both in the type and number of chemicals biotransformed, ranging from 2 to 24 and 4 to 27 biotransformed mutagens per bacterial species under anaerobic and microaerobic conditions, respectively (Fig. 2C-D). Notably, a fraction of tested compounds was more strongly metabolized by anaerobes belonging to the genera *Bacteroides, Collinsella, Parvimonas, and Solobacterium* compared to anaerobes from other genera such as *Veillonella* and to most of the microaerobes. Such metabolic differences could not be explained by differences in biomass (measured as OD600) of cultures from different bacterial species, as a range of culture densities was measured among the strong metabolizers (Fig. 2C and Table S2). Further, we observed only a poor correlation between the number of chemicals biotransformed and biomasses of the corresponding bacterial cultures (Fig. S1A). These observations indicate that there are interspecies differences in the metabolic potential to biotransform organic mutagens.

**Figure 2.**
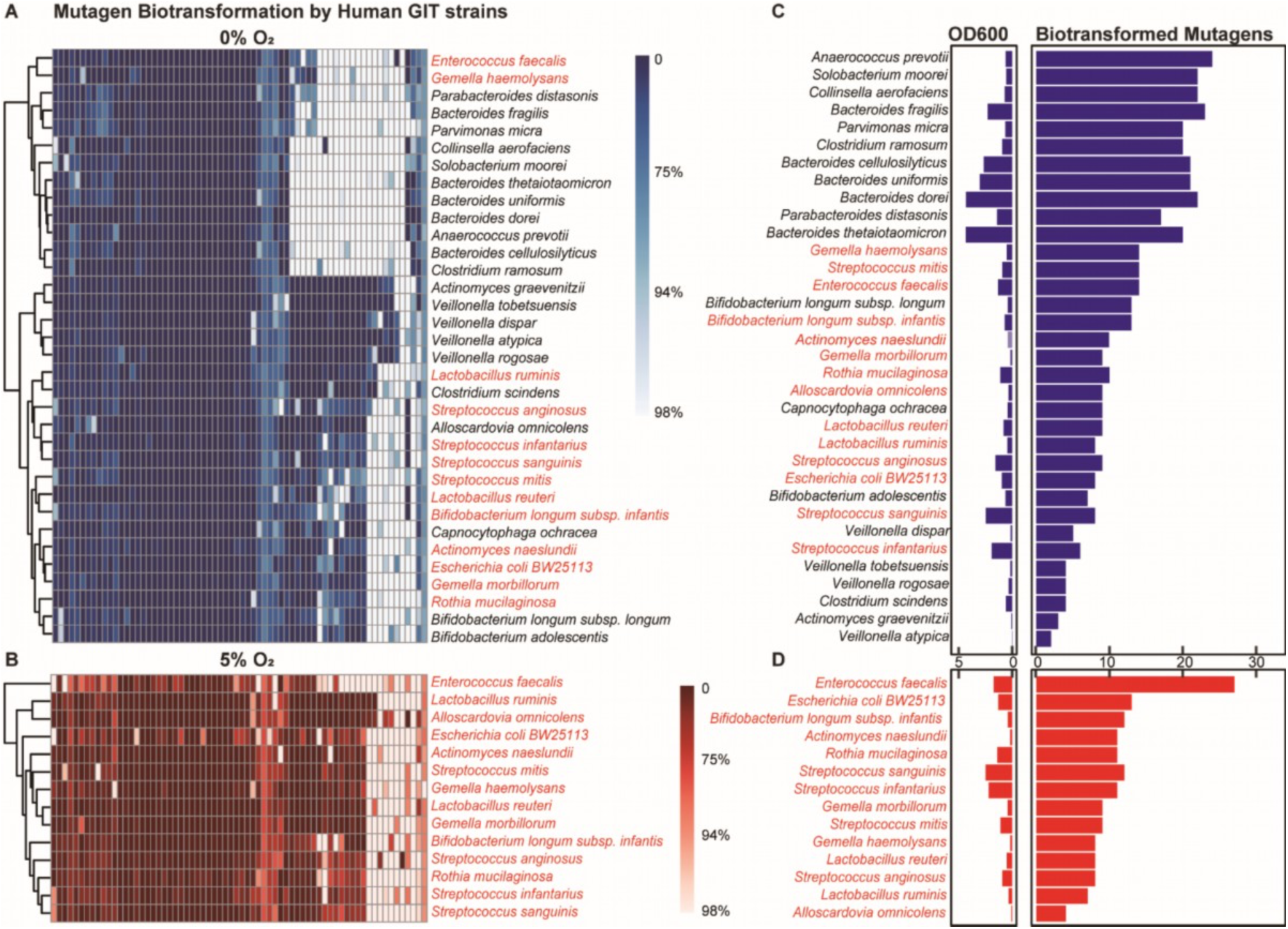
Biotransformation of organic carcinogens by human gut bacteria under anaerobic and microaerobic conditions. **A-B.** Average percentage (n = 4) of individual mutagens metabolized by 34 human gut bacterial species under anaerobic (**A**: 0% O_2_, blue heatmap) and microaerobic (**B**: 5% O_2_, red heatmap) conditions. Names of non-obligate anaerobe bacteria are shown in red. **C-D.** Number of carcinogens metabolized (>75%, adj. p-value = 0.05) by each species under anaerobic (C) and microaerobic (D) conditions and bacterial culture density (measured as OD600) of the respective bacterial culture after 12 hours of incubation.

To gain mechanistic insights into why certain mutagens were more potently metabolized than others, we performed an overrepresentation analysis of the chemical groups present in compounds that are either biotransformed and non-biotransformed by bacteria (Fisher’s exact test p-values < 0.05, Fig. S1B-C and Table S4). This analysis revealed that nitro and azo groups rendered compounds more prone to bacterial biotransformation under anaerobic conditions but not under microaerobic conditions (Fig. S1B-C), which can be explained by bacterial reduction reactions occurring under anaerobic conditions. This is further corroborated by the observation that nitro- and azo-containing compounds were most strongly metabolized by anaerobic species and demonstrated the highest interspecies variation in the extent of biotransformation (Fig. 2E and Table S5-6).

Together, these results demonstrate that around two-thirds of the tested mutagens can be biotransformed by human gut bacteria with pronounced inter-species differences. Notably, in addition to the better studied bacterial isolates from the large intestine, we also found species isolated from the oral cavity and small intestine such as *Anaerococcus prevotii* and *Solobacterium moorei* to be strong mutagen metabolizers.

### Bacterial metabolism alters activities of mutagens in a species-dependent manner

The high metabolic capacity of human gut bacteria against a diverse set of mutagens and the observed inter-species differences in mutagen biotransformation led to the question about the functional consequences of bacterial mutagen biotransformation. To address this question, we directly measured the impact of bacterial biotransformation on mutagen DNA reactivity by establishing a pipeline to couple bacterial biotransformation assays to Ames reversion-based mutagenicity tests. This allowed us to simultaneously monitor the bacterial biotransformation of mutagens and changes in their mutagenic potential upon biotransformation (Fig. 3A).

**Figure 3.**
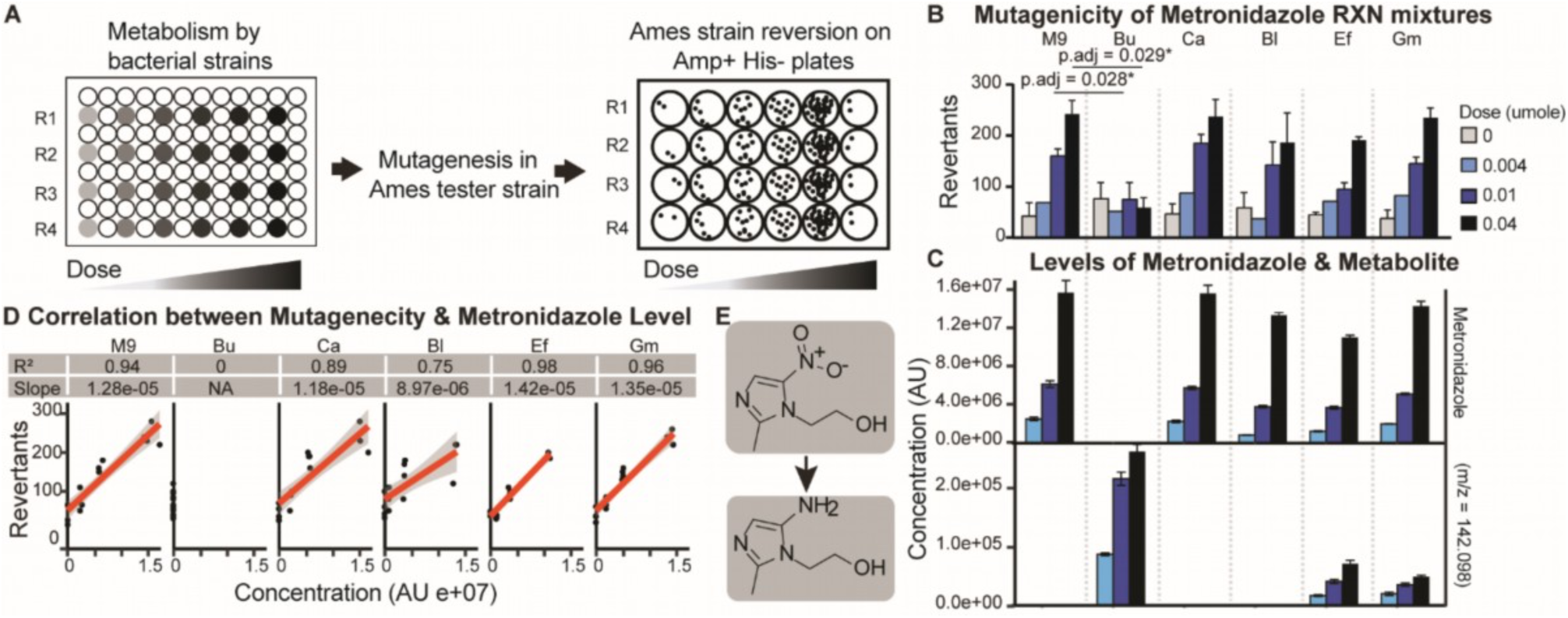
Mutagenicity alterations and biotransformation of metronidazole by bacteria. **A.** Scheme of the experimental setup to couple Ames mutagenesis tests to gut bacterial biotransformation assays at a range carcinogen concentrations to test mutagenicity changes following bacterial metabolism of mutagens with four replicates (‘R1’-‘R4’). **B.** Mutagenicity of 12-hour reaction mixtures (n = 4) of metronidazole incubated with different gut bacterial species in Ames tester species TA100 for a dose range of 0-0.04 umole. Adjusted p-values shown for dose 0.01 and 0.04 umole between M9 and *B. uniformis*. Standard deviations from the means of each group shown as error bars. Pairwise comparison between Bu and M9 reaction mixtures at respective concentrations performed with Wilcoxon test and p-values were adjusted with Bonferroni correction. **C.** Levels of metronidazole and its amino derivative ([M+H] m/z 142.098) measured in reaction mixtures after 12 hours of bacterial incubation. Standard deviations from the means of each group shown as error bars. **D.** Correlation between the number of revertants of each reaction mixtures and the metronidazole levels after 12 hours of bacterial incubation. Significance and slope of correlations for each species was calculated with Pearson’s correlation test and indicated in respective graphs. **E.** Chemical structures of metronidazole and its amino derivative ([M+H] m/z 142.098). M9 = M9 media without bacteria as negative control, Bu = *B. uniformis*, Ca = *C. aerofaciens*, Bl = *B. longum*, Ef = *E. faecalis*, Gm = *G. morbillorum*.

To assess the link between bacterial mutagen biotransformation and altered mutagenicity, we first used the direct mutagen metronidazole^18^, which is an antimicrobial agent allowing us to titrate the dose to assess mutagenicity rather than bacterial killing. Metronidazole was found to be efficiently metabolized by *Bacteroides uniformis* but not by *Collinsella aerofaciens, Bifidobacterium longum subsp. longum, Enterococcus faecalis*, and *Gemella morbillorum* in our initial screen (Fig. 2A). We incubated these bacteria with metronidazole within its mutagenic concentration range in the Ames test (0-2 mM), monitored biotransformation reactions (*i.e.*, parent compound and its biotransformation products) by LC-MS and LC-MS/MS. We then exposed an Ames tester strain (*Salmonella typhimurium* TA100) with each of the biotransformation culture supernatants for 12 hours to then measure mutagenicity by counting revertant colonies following plating of the Ames tester strain. We found that both the levels and the mutagenicity of metronidazole were only reduced after incubation with *B. uniformis* (Fig. 3B-C and Table S7). Indeed, a positive correlation was observed between the metronidazole level in each reaction mixture of each gut bacterial species with the mutagenicity of its respective reaction mixture with the exception of *B. uniformis* (Fig. 3D and Table S8). This suggested that bacterial biotransformation of metronidazole reduces its mutagenic potential. Using untargeted metabolomics analysis, we found that the reductive amine-metabolite with the mass 141.090 Da is indeed most strongly produced by *B. uniformis* (Fig.3C, Fig. S2A and Table S15). Moreover, the reduction of the nitro group of metronidazole to an amine has been described previously to deactivate metronidazole genotoxicity^19^, which corroborates our hypothesis that the nitro-reductive metabolism of metronidazole by *B. uniformis* is responsible for the reduction of its mutagenicity. These results illustrate the use of our developed experimental pipeline, coupling bacterial carcinogen biotransformation with mutagenicity assays to systematically test the impact of bacterial metabolism on the activity of organic carcinogens.

We next asked whether bacterial biotransformation of mutagens would always lead to reduced mutagenicity. To test this, we selected another strong mutagenic carcinogen^20^, 4-nitroquinoline-1-oxide (4NQO), which can covalently bind to DNA molecules^21^ and induce mutagenesis without prior enzymatic bioactivation. 4NQO, containing a nitro group, was strongly metabolized by 29 species in our initial screen (Fig. 2C). Intriguingly, unlike metronidazole, biotransformation of 4NQO did not always lead to reduced mutagenicity (Fig. 4A and Table S9), which was only the case for two out of the four metabolizing bacterial species tested. In fact, mutagenicity of the complete Ames-sensitive dose range was reduced by *B. uniformis* and *C. aerofaciens*, but remained unchanged following incubation with *B. longum* and *G. morbillorum*. To better understand this observation, we asked whether the metabolic products generated by these bacteria and their mutagenicity might be different. Using untargeted metabolomics measurements, we identified three different bacterial biotransformation products of 4NQO (with monoisotopic masses 176.059 Da, 160.064 Da and 144.069 Da) accumulating at different levels following incubation with each of the four gut bacterial species (Fig. 4B, Fig. S3A and Table S16). Tandem mass spectrometry analysis identified these 4NQO metabolites as biotransformation products resulting from the reduction of both the nitro and the nitroxyl group (Fig. 4C). Among these biotransformation products, the first nitro-reduction intermediate, 4-hydroxyaminoquinoline-1-oxide (4HAQO, 176.059 Da), was strongly accumulating in *B. longum* and *G. morbillorum* cultures (Fig. 4B and Fig. S3B), while *B. uniformis* and *C. aerofaciens* further reduced 4HAQO to amino derivatives with monoisotopic masses of 160.064 and 144.069 Da (Fig. 4C). Together with the mutagenicity rates observed for the different bacterial species, these results suggested that the *N*-hydroxyl metabolite accumulating in *B. longum* and *G. morbillorum* reaction mixtures has comparable mutagenic activity as the parent compound 4NQO. Indeed, 4HAQO was previously shown to interact with DNA^21,22^. To quantify these findings, we assessed 4HAQO dose dependency of the mutagenicity of *B. longum* and *G. morbillorum* reaction mixtures and found a comparable correlation for each of the two species (Fig. 4D and Table S10). To further investigate the strain-specific accumulation of 4HAQO, we performed a time-resolved biotransformation assay with 4HAQO as a pure chemical. We found that the conversion of 4HAQO to its amino derivatives (*i.e.*, 160.064 Da and 144.069 Da) followed a much slower kinetics in *B. longum* and *G. morbilorum* compared to *B. uniformis* cultures (Fig. 4E and Table S11). Together, these results indicate that biotransformation of the same carcinogenic compound by different bacterial species can result in the accumulation of different biotransformation products that can have marked differences in their mutagenic potential.

**Figure 4.**
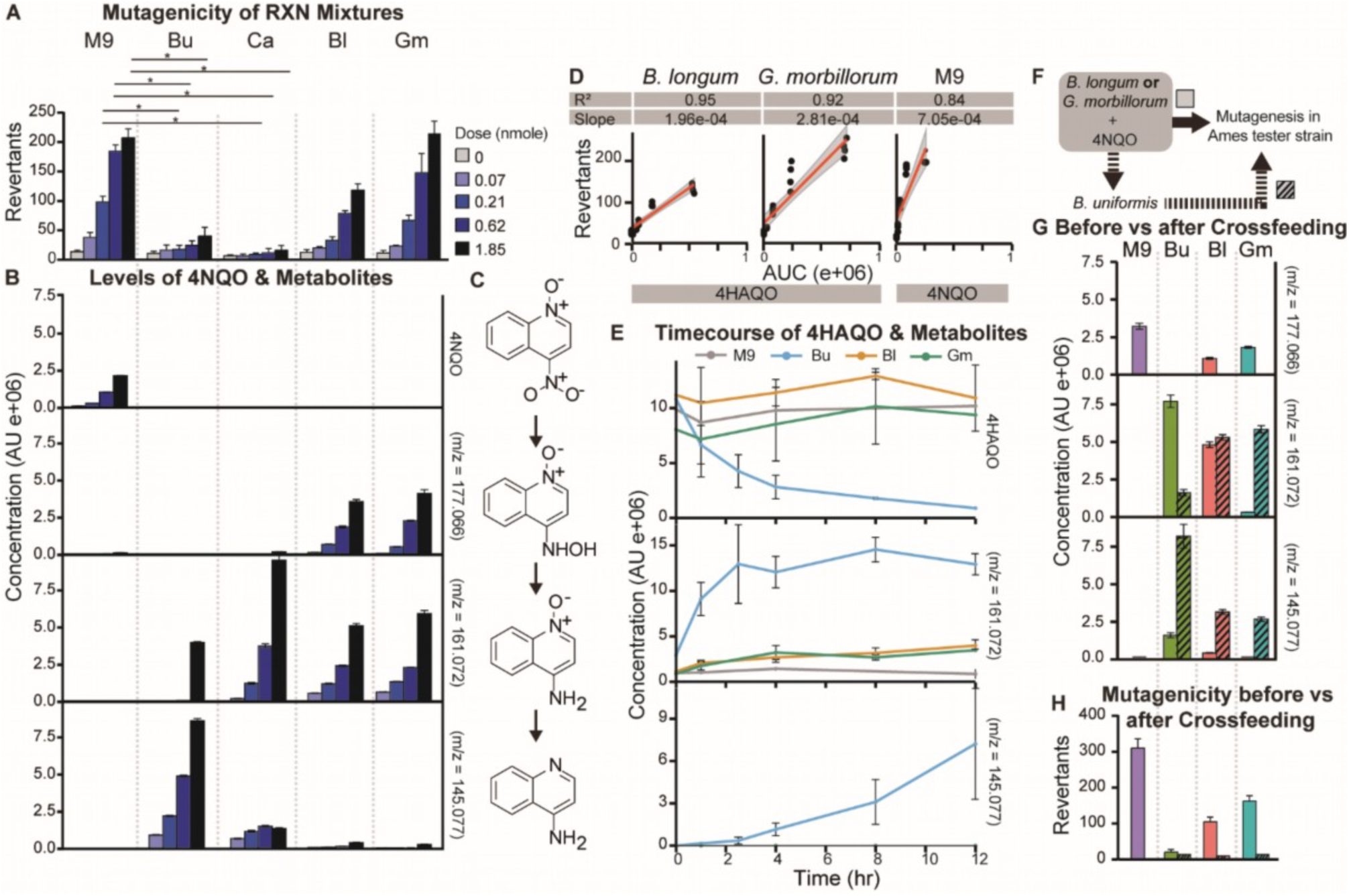
Inter-species differences in 4-nitroquinoline-1-oxide (4NQO) biotransformation and changes in mutagenicity. **A.** Mutagenicity of 12-hour reaction mixtures (n = 4) of 4NQO incubated with different bacterial species in Ames tester species TA100 for a dose range of 0 to 1.85 nmole. Standard deviations from the means of each group shown as error bars. Pair-wise comparison between Bu/Ca reaction mixtures and M9 reaction mixtures at respective concentrations performed with Wilcoxon test and p-values were adjusted with Bonferroni correction. *, p-value < 0.05. **B**. Quantification of 4NQO (area under the curve/AUC) and its reduced metabolites with [M+H] m/z 177.066, 161.072 and 145.077 in reaction mixtures (n = 4) after 12 hours of gut bacterial incubation. Standard deviations from the means of each group shown as error bars. **C.** Biotransformation pathway of 4NQO and reaction intermediates. **D.** Correlation between the number of revertants of the reaction mixtures of *B. longum* and *G. morbillorum* and the first metabolic intermediate with m/z 177.066 (4HAQO) and between the number of revertants in M9 and the parent compound 4NQO. Significance and slope of correlations for each species was calculated with Pearson’s correlation test and indicated in respective graphs. **E.** Biotransformation kinetics of 4HAQO (n = 4) over 12 hours of incubation with *B. uniformis, B. longum*, and *G. morbillorum.* **F.** Experimental setup to test cross-feeding of carcinogen metabolites from reaction mixtures of 4NQO with *B. longum* or *G. morbillorum* with *B. uniformis* for further metabolism and mutagenicity assessment. **G.** Abundance of 4HAQO and its amino derivatives measured from reaction mixtures of either supernatants of 12-hour reaction mixtures (n = 4) of 0.62 nmole 4NQO with *B. longum* and *G. morbillorum* (without dashes, as in the original setup) or from the 12-hour reaction mixtures of these supernatants cross-fed to *B. uniformis* (dashed, as in **F**). Standard deviations from the means of each group shown as error bars. **H.** Mutagenicity of the respective reaction mixtures in **G** tested in TA100. M9 = M9 media without bacteria, Bu = *B. uniformis*, Ca = *C. aerofaciens*, Bl = *B. longum*, Gm = *G. morbillorum*.

Given that different bacterial species exist in microbial gut communities, we wanted to investigate whether metabolic cross-feeding from non-detoxifying species to detoxifying species could lead to detoxification. To this aim, we tested whether the mutagenic biotransformation product of 4NQO, 4HAQO, produced by *B. longum* and *G. morbillorum* could be further metabolized and detoxified by *B. uniformis* (Fig. 4F). Indeed, we found that 4HAQO produced by *B. longum* and *G. morbillorum* is completely metabolized by *B. uniformis* after transferring their 4NQO-incubated culture supernatant to *B. uniformis* cultures (Fig. 4G and Table S12), and that mutagenicity was reduced accordingly (Fig. 4H and Table S13).

Altogether, using metronidazole and 4NQO as example mutagens, we demonstrated that species-specific differences in bacterial carcinogen biotransformation activity exist, which can lead to alteration of the mutagenic potential of carcinogens. We further showed that cross-feeding mechanisms between gut bacteria can modulate carcinogen biotransformation and resulting mutagenicity. This demonstrates the possibility to harness such cross-feeding mechanisms for targeted bacterial detoxification of carcinogens.

### Interactions between host and gut bacterial biotransformation alters overall mutagenicity of procarcinogens

A large number of organic mutagens require activation by human enzymes to become mutagenic, known as bioactivation. Bioactivated procarcinogens produced by the liver could be further metabolized in the gut following biliary excretion and vice versa, microbially produced procarcinogen metabolites in the gut could be bioactivated by liver enzymes following intestinal absorption. For these reasons, we asked how the biotransformation activities of gut bacteria and its human host interact and how such interactions can impact the overall mutagenicity of a procarcinogen. To this aim, we further extended our experimental workflow to combine bacterial mutagen biotransformation with human liver metabolism and Ames mutagenicity assays (Fig. 5A).

**Figure 5.**
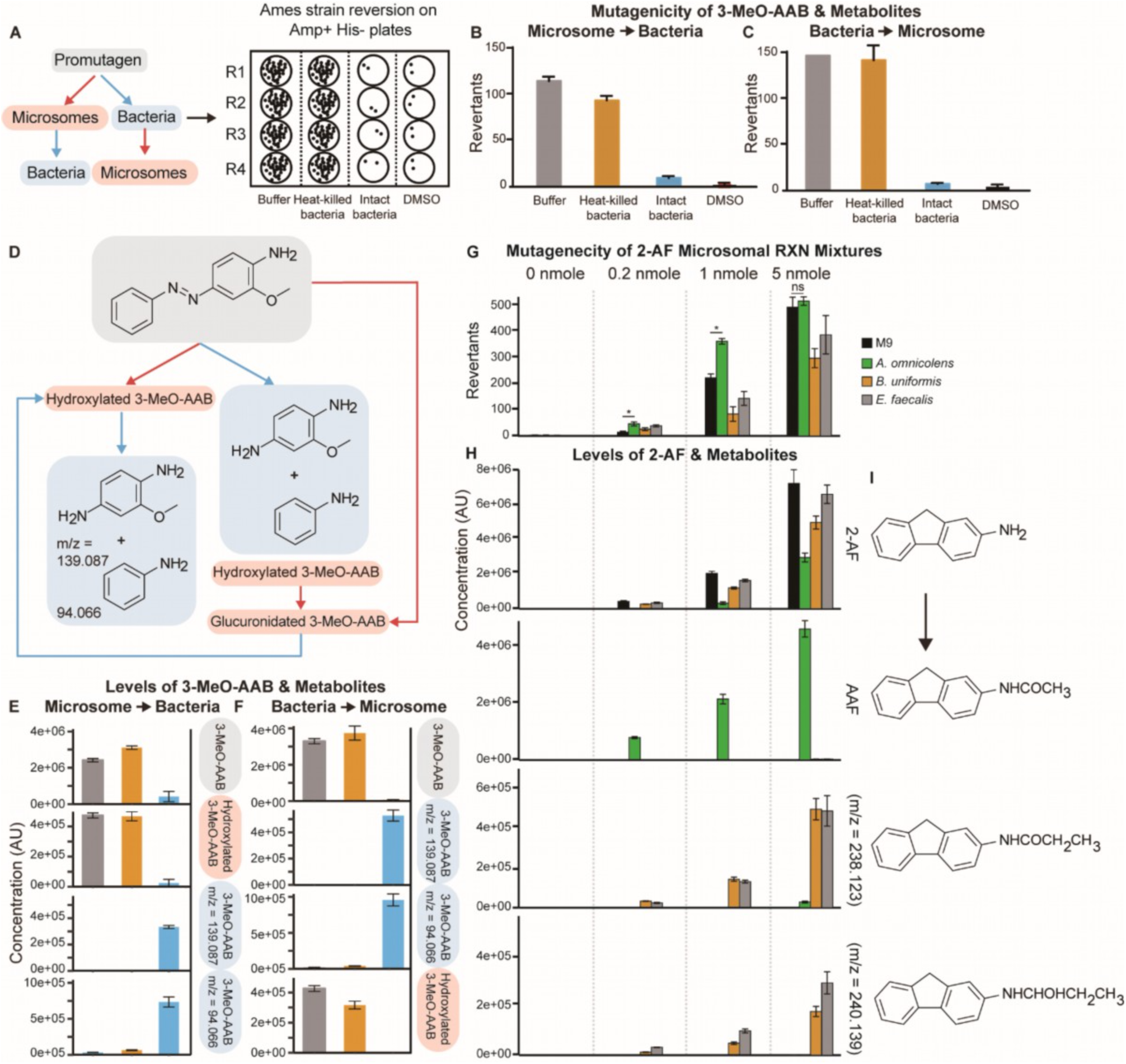
Metabolic bacteria-liver interactions and mutagenicity alterations of 3-methoxy-4-aminoazobenzene (3-MeO-AAB) and 2-aminofluorene (2-AF). **A.** Experimental setup to test metabolic interactions of procarcinogens between gut bacteria and liver enzymes. In this *in vitro* setup, the promutagen is first incubated with bacteria or human microsomes in four replicates (‘R1’-‘R4’), respectively, followed by reaction mixture transfer to liver microsome or gut bacteria, before the overall mutagenicity is assessed using Ames reversion assays. **B-C.** Mutagenicity of 3-MeO-AAB reaction mixtures in Ames tester strain TA98 after a 4-hour gut bacterial biotransformation (n = 4) of microsome bioactivated procarcinogens (**B**) and microsomal reaction mixture (n = 4) of gut bacterial biotransformation assays (**C**). Standard deviations from the means of each group shown as error bars. **D.** Carcinogen metabolites resulting from microsomal (host associated) enzymatic reactions shown in red and resulting from bacterial biotransformation reactions shown in blue. **E-F.** LC-MS quantification of 3-MeO-AAB and its metabolites following the reaction orders in **B** (**E**) and in **C** (**F**) (n = 4). **G.** Mutagenicity of the supernatants of 12-hour reaction mixtures of 2-AF incubated with different bacterial species followed by human microsome bioactivation for 4 hours within a dose range of 0-5 nmole in Ames tester strain TA98. Pair-wise comparison between Ao reaction mixtures and M9 reaction mixtures at respective concentrations performed with Wilcoxon test and p-values were adjusted with Bonferroni correction. P-value cutoff: * = 0.05. **H**. LC-MS quantification of 2-AF (area under the curve/AUC) and its metabolites acetylaminofluorene (AAF) and other putative metabolites with [M+H] m/z 238.123 and 240.139 in reaction mixtures after 12 hours of bacterial incubation. **I.** Structures of 2-AF biotransformation products.

We selected the procarcinogenic azo dye 3-methoxy-4-aminoazobenzene (3-MeO-AAB), which has been reported to require *N-*hydroxylation by cytochrome P450 to induce mutagenicity^23^. Indeed, when we incubated the chemical with human microsomes, 3-MeO-AAB markedly induced mutagenesis in the Ames tester strain (*i.e.*, *S. typhimurium* TA98) (Fig. 5B, column ‘Buffer’ and Table S14). To simulate biliary excretion *in vitro*, we transferred human microsome reaction mixtures containing bioactivated 3-MeO-AAB to *B. uniformis* cultures and heat-killed control cultures to assess the mutagenic potential of the liver-bioactivated carcinogen after additional gut bacterial biotransformation. Following gut bacterial biotransformation, but not the control incubation containing heat-inactivated bacteria, the mutagenic activities of the microsomally activated carcinogen metabolites were almost completely diminished (Fig. 5B and Table S14). Using untargeted metabolomics analysis, we found that the metabolite produced from human microsomal reaction mixture was the expected hydroxyl-3-MeO-AAB (monoisotopic mass 243.101 Da, Fig. S4A, Table S17 and Fig. 5D-E). Intriguingly, we only detected the azo-reduced metabolites with isotopic masses 138.079 and 93.058 Da from the sequential bacterial reaction mixtures (Fig. S4B, Table S19, Fig. 5D-E and Table S20), possibly because the *N-*hydroxylated 3-MeO-AAB followed first an azo then an *N-*hydroxyl reduction. Overall, these results demonstrate that *B. uniformis* can detoxify the mutagenic metabolite of 3-MeO-AAB produced by human liver enzymes, likely followed by biliary elimination to reach the intestine.

Next, we asked whether the promutagen could already be inactivated by gut bacteria prior to bioactivation by host enzymes. To test this, we first subjected 3-MeO-AAB to biotransformation by *B. uniformis* under anaerobic conditions. We then exposed the supernatant of this bacterial reaction mixture to human liver metabolism employing microsomal preparations. We found that human microsomes could not further activate the gut bacterial biotransformation product of 3-MeO-AAB (in contrast to heat-killed bacterial culture controls and buffer controls) to induce mutagenicity in the subsequent Ames tests (Fig. 5C and Table S14). Using untargeted metabolomics analysis, we found that *B. uniformis* efficiently reduces 3-MeO-AAB to its azo-reduced metabolites with monoisotopic masses 138.079 and 93.058 Da (Fig. S4C, Fig. 5F, Tables S18 and S20), which we found to not be further metabolized and activated by human microsomes (Fig. 5C). These results suggest that the promutagen 3-MeO-AAB is efficiently detoxified by *B. uniformis* preventing it from subsequent bio-activation by human enzymes.

Altogether, our findings illustrate how human gut bacteria can impact the mutagenic activity of pro-carcinogens both through biotransformation and inactivation of promutagens that have been bioactivated by human enzymes and through biotransformation and inactivation of promutagens that can then no longer be bioactivated by human enzymes.

### Bacterial metabolism alters activities of promutagens in a species-dependent manner

As we found pronounced inter-species differences in gut bacterial carcinogen biotransformation (Fig. 2), we wondered how these differences would affect procarcinogen interactions between gut bacteria and the host. To this aim, we tested the metabolism of the synthetic promutagen, 2-aminofluorene (2-AF), in three different gut bacterial species, *A. omnicolens, B. uniformis* and *E. faecalis* under anaerobic conditions. Although all three bacterial species were able to biotransform 2-AF to various extents, we only observed a reduction in mutagenicity following the incubation with *B. uniformis* and *E. faecalis* prior to microsomal activation (Fig. 5G-H and Table S21). In contrast, biotransformation of 2-AF by *A. omnicolens* even increased the mutagenicity by 3.2-fold at 0.2 nmole (two-tailed test, p-value = 0.001, n = 4) and by 1.6-fold at 1 nmole (two-tailed test, p-value = 9.05e-6, n = 4) (Fig. 5G and Table S21). To explain this observation, we quantified the relative amounts of the most prominent metabolite candidates of 2-AF and found that the acetylated metabolite of 2-AF, *N*-acetyl-2-aminofluorene (AAF) was produced extensively by *A. omnicolens* but not by the other strains (Fig. 5H and Table S22). In contrast, two different 2-AF metabolites with m/z values 238.123 and 240.139 were produced in high amounts by the other two bacteria tested, but not *A. omnicolens* (Fig. 5H and Table S22). These results likely explain the observed differences in mutagenicity upon 2-AF incubation with gut bacteria, as liver bioactivation requires the formation of a nitrenium ion, for which AAF is a more efficient substrate than 2AF (and likely also the additional gut bacterial 2AF metabolites identified). Together, these results demonstrate that the observed species-differences in gut bacterial promutagen metabolism can lead to the generation of different procarcinogen metabolites with modified (increased and decreased) bioactivation potential by host enzymes, resulting in gut-microbiome dependent mutagenicity.

### In vivo validation of microbiota-host interplay in procarcinogen metabolism

To *in vivo* validate our *in vitro* findings demonstrating the interplay between gut bacterial and human enzyme systems modulating procarcinogen mutagenicity (Fig. 4), we established a gnotobiotic mouse model (Fig. 6A). We stably colonized germ-free C57BL/6 mice with *B. uniformis*, administered the procarcinogen 3-MeO-AAB in drinking water for 7 days to then monitor steady-state levels of the procarcinogen and its bioactivated metabolite, hydroxyl-3-MeO-AAB, in different body compartments of colonized and germfree control animals. While the levels of 3-MeO-AAB were comparable in the small intestine between the two groups of animals, the levels decreased along the intestinal tract only in mice colonized with procarcinogen-metabolizing bacteria (Fig. 6B and Table S23-24). This data demonstrates that the ingested procarcinogen is not fully absorbed in the small intestine and reaches the lower intestine where gut bacteria can biotransform it to its non-bioactivatable form. Intriguingly, we only detected low levels of intestinal hydroxyl-3-MeO-AAB, the bioactivated procarcinogen resulting from phase I metabolism in the liver, in either group of mice (Fig. 6C and Table S23-24). This suggests that the bioactivated procarcinogen is only poorly eliminated through biliary excretion and that glucuronidation (phase II metabolism) might be required to transport the hepatically activated procarcinogen back into the intestinal tract. Indeed, we found comparable levels of glucuronidated hydroxyl-3-MeO-AAB (m/z 418.125 [M-H^+^]) in the small intestine of both mouse groups, suggesting similar levels of host enzymatic activities independent of the presence of gut colonization (Fig. 6D and Table S23-24). However, upon further propagation in the intestinal tract, the levels of the glucuronidated metabolite were strongly decreasing in colonized mice only (Fig. 6D). This can be explained by the presence of gut bacterial glucuronidases that readily deconjugate the metabolite to regenerate the hydroxylated and mutagenic form of 3-MeO-AAB. The bioactivated mutagen is then further subjected to bacterial azo-reduction leading to its gut bacterial deactivation, as demonstrated in our *in vitro* assay (Fig. 5D and Table S23-25). To further corroborate this observation, we performed *in vitro* deglucuronidation of the gastrointestinal samples demonstrating that the bioactivated procarcinogen, hydroxyl-3-MeO-AAB, indeed enters the gut as a glucuronide conjugate and that its inactivating biotransformation depends on the presence of gut bacteria (Fig. 6E-F and Table S23-25). Intriguingly, we did not detect the bacterial biotransformation products of 3-MeO-AAB and hydroxyl-3-MeO-AAB *in vivo*, possibly due to further metabolism and/or absorption. Altogether, this gnotobiotic mouse model validates our *in vitro* finding of a tight interaction between gut bacterial and host procarcinogen metabolism.

**Figure 6.**
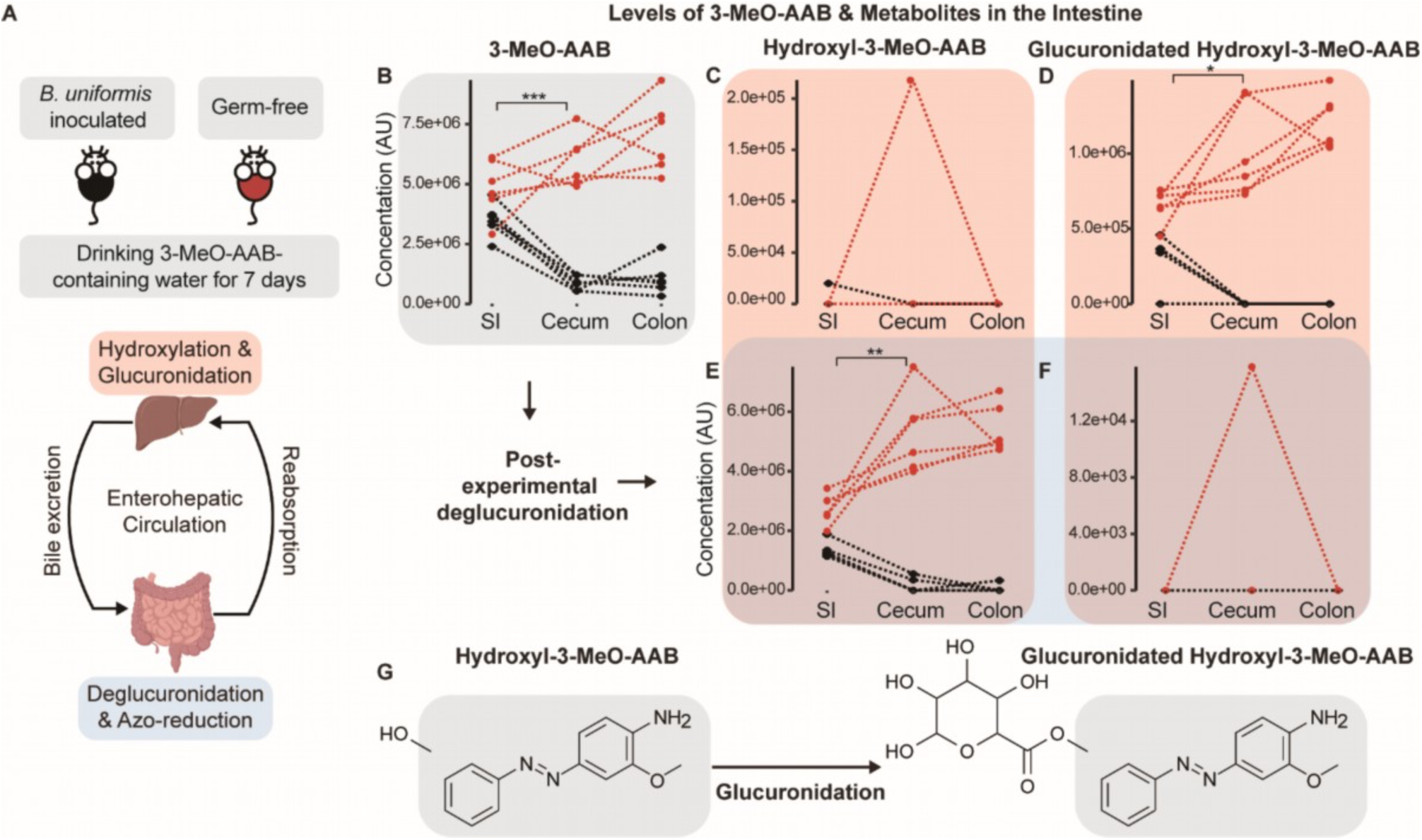
Gnotobiotic mouse model shows gut bacteria-host interaction in procarcinogen metabolism. **A.** Experimental design for testing host-microbe interactions in 3-MeO-AAB metabolism in gnotobiotic mice and illustration of *in vivo* metabolic pathways. **B-F.** Concentration of 3-MeO-AAB, its host-derived hydroxylation metabolite and glucuronidated metabolite of the hydroxyl-3-MeO-AAB in the intestine of *B. uniformis-*inoculated mice (black) and germ-free mice (red), before (**C-D**) and after (**E-F**) post-experimental deglucuronidation, n = 6/6. Pairwise comparisons between tissues within the germ-free group were done with estimated marginal means (EMMs) after fitting the data with a linear mixed model, and p-values were adjusted for multiple testing. P-value cut-offs: 0.001 = ***, 0.01 = **, 0.05 = *. **G.** Scheme for glucuronidation reaction of the hydroxylated metabolite of 3-MeO-AAB.

## Conclusions

The human microbiome and environmental organic carcinogens, as part of the human exposome, have both been associated with tumorigenesis and cancer development. In this study, we aimed at establishing a new causal link between the human gut microbiome and organic mutagens through systematic mapping of the capacity of gut bacteria to biotransform carcinogens. We found that the majority of tested carcinogens (41 of 68) are indeed metabolized by at least one member of a panel of 34 bacterial species, representative for the upper and lower human intestinal tract. Further, we observed pronounced interspecies variation in the metabolic potential of these bacteria to metabolize carcinogens. To gain insights into the functional consequences of the observed gut bacterial carcinogen metabolism, we developed an *in vitro* pipeline coupling biotransformation assays with mutagenicity testing. This pipeline allowed both the monitoring of mutagenicity changes accompanying bacterial biotransformation and the testing of the impact of gut bacterial cross-feeding and microbiota-host interactions on carcinogen mutagenicity. To validate our *in vitro* findings *in vivo*, we set up a gnotobiotic mouse model to demonstrate that the metabolic crosstalk between gut bacteria and their host leads to altered procarcinogen activation.

Metabolic interactions between gut bacteria and between gut bacteria and their host are often complex. Our modular *in vitro* approach enables to study steps of these interactions individually and in combination, such as the biotransformation by different bacteria and liver enzymes. For example, bacterial cross-feeding is well-known for primary metabolites leading to mutualistic interactions between bacteria^24^. Here, we demonstrate that cross-feeding of genotoxin-derived metabolites between bacteria may benefit the host (and the microbiome) when genotoxic metabolites generated from one bacterial species can be detoxified by another.

We focused our study on DNA-reactive, organic carcinogens. However, genotoxicity can also be induced by inorganic compounds via a variety of other pathways that do not directly target DNA, but which induce inflammation and the generation reactive oxygen species (ROS) attacking DNA. Furthermore, tumorigenesis is a multifactorial process, to which other factors than chemicals contribute and which we do not capture in our models. Our developed pipeline can however be modified to investigate additional aspects of the interaction between carcinogens, the microbiome and human cells. For example, the bacterial and liver biotransformation assays could be coupled to alternative high-throughput genotoxicity readouts, such as the comet assay for a broader coverage of genotoxicity mechanisms.

With our work, we aimed at showcasing the metabolic potential encoded by gut bacteria to metabolize organic mutagens and the resulting functional consequences. Overall, our study emphasizes the role of a class of enzymes that has been largely overlooked in traditional genotoxicity studies, the gut bacterial enzymes. Such additional metabolism in the host might significantly alter the systemic exposure of carcinogens and as a consequence, tumorigenesis and cancer development. Our developed pipeline, the generated data, and our findings provide a foundation and resource for future studies that aim at investigating the functional role of gut microbes in cancer onset and development and that assess the gut microbiome as potential predisposition risk for carcinogen exposure.

## Methods

### Chemicals

68 (pro)mutagens were purchased from Sigma and Ambeed (Table S1). All chemicals were dissolved in DMSO and stock solutions of 10 mM were stored at −70°C. 40 combinatorial pools were prepared for these chemicals with the script from https://git.embl.de/bartmans/PoolingScheme.

### Bacterial species

All bacterial species used in this study and their sources are listed in Table S2.

### Bacteria cultivation and biotransformation assays

Frozen glycerol stocks of bacterial species were streaked on pre-reduced modified Gifu Anaerobic Medium (mGAM) agar plates (HyServe GmbH & Co. KG) and incubated for 24-48 hours at 37°C until single colonies could be picked, in an anaerobic chamber (Coy Laboratory Products, containing 3% H_2_, 12% CO_2_, and 75% N_2_). Four colonies of each species were picked, inoculated in two 10 mL-round bottom tubes (Greiner) of mGAM medium, incubated at 37°C overnight under anaerobic and microaerobic (Coy Laboratory Products, containing 5% O_2_, 5% CO_2_, and 90% N_2_) conditions, shaken on an orbital tube shaker (neoMix, Art. No. 7-0921) at 500 rpm, respectively. Bacterial growth was monitored by OD600 measurements. Bacterial cultures between OD 0.09-5.00 were centrifuged at 3000 rpm for 10 minutes and the supernatants were discarded in the growth chamber. To preserve sufficient biomass for efficient biotransformation and prevent overgrowth of the cultures, 10 mL of M9 minimal medium (49.3 mM Na_2_HPO_4_, 22 mM KH_2_PO_4_, 18.8 mM NH_4_Cl, 8.6 mM NaCl, 100 uM CaCl_2_, 2 mM MgSO_4_, 0.4% glucose) was used to resuspend the original cultures. 200 uL of resuspended cultures of each species was added to each of the 40 pools containing 4 uL of the chemical mixture and into 4 control wells containing DMSO. The final concentration of each chemical assayed was 14.3 uM and 2% DMSO. After 0, 4 and 12 hours of incubation under anaerobic or microaerobic conditions on a plate shaker (BioSan, PST-100HL), reaction mixtures were mixed thoroughly with a 100 uL multichannel pipette, and 10 uL of the reaction mixture were transferred to V-bottom 96-well storage plates (Fisher Scientific, Cat. No. 10304513), snap-frozen on dry ice, and stored at −70°C until further processing.

### Ames tester species preparation

Tester species *S. typhimurium* TA98 and TA100 could be reverted by most of the tested mutagens according to previous literature and were hence used as the main tester species in the current study. Strains were first checked for genetic integrity as previously described^25^. Glycerol stocks were streaked on enriched glucose minimal plates (3.3 uM biotin, 257.8 uM histidine, 0.024 mg/ml ampicillin) and incubated aerobically at 37°C. A single colony was picked for each tester species and inoculated in 10 mL nutrient broth medium (Thermo Scientific, OXCM0067B) at 37°C, 200 rpm for 12-14 hours. Prior to the Ames test, the cultures were 10-fold concentrated in phosphate buffer (100 mM, pH 7.4) after being washed twice.

### Ames test for direct mutagens following bacterial metabolism

Mutagens were tested as pure chemicals against Ames tester species TA98 or TA100 in the dose range of 0-1 umole with eleven 3-fold serial dilutions. The doses that elicited dose-dependent mutagenesis without inducing toxicity were selected as the testing dose range as previously reported^25^. To test the effect of bacterial metabolism on chemical mutagenicity, mutagens in their respective non-toxic, mutagenic dose ranges were incubated with bacterial culture that has been resuspended and 10-fold concentrated in M9 minimal medium, as well as boiled culture (for 30 min) or M9 minimal medium as controls. After incubation / bacterial biotransformation, 25 uL of the reaction mix were pre-incubated for 1 hour at 37°C, 500 rpm on a plate shaker (BioSan, PST-100HL) with 5 uL of phosphate-buffer-washed and 10-fold concentrated Ames tester species culture. 100 uL melt 0.35% top agar (6 mg/mL NaCl, 50 uM biotin/histidine solution, 0.35% agar) was then mixed with each reaction mix and transferred to wells in 24-well glucose-minimal plates. After 48 hours of aerobic incubation at 37°C, colonies were counted.

### Ames test for promutagens after metabolism by human microsomes and strict anaerobes

Promutagens were first activated by incubating in equal volumes with a 2-fold human microsomal (ThermoFisher Cat. No. HMMCPL) reaction mixture containing 1mg/mL microsomes, 0.8 units/mL glucose-6-phosphate dehydrogenase, 2 mM of NADP, NADH and NADPH and 5.46 mM glucose-6-phosphate in 100 mM phosphate buffer. Alternatively, promutagens were metabolized by incubating with equal volumes of 10-fold concentrated bacterial culture. After incubation, 20 uL of the reaction mixture were metabolized by 20 uL of 10-fold concentrated bacterial culture or alternatively, 2-fold concentrated microsome reaction mixture as described above for another 1.5 hours. The reaction mixture was then preincubated with 10 uL of 10-fold concentrated Ames tester species for 1 hour and mixed with 100 uL top agar before being transferred to glucose minimal agar, as described above.

### Ames test for promutagens after metabolism by human microsomes and facultative anaerobes

Prior to Ames tests, the metabolizer species were streaked onto the glucose minimal plates containing ampicillin to check for background growth of the metabolizer facultative species which could interfere with the Ames species reversion. In case of no background growth of the microaerobic metabolizer species, experiments were performed as described for strict anaerobes above.

### Sample preparation for LC-MS measurements

Solid tissues and liquid samples were prepared for LC-MS and LC-MS/MS analysis by organic solvent extraction (acetonitrile:methanol, 1:1) at −20 °C after the addition of internal standard mix (sulfamethoxazole, caffeine, ipriflavone and yohimbine each to a final concentration of 80 nM) as previously described^7^.

### LC-MS and LC-MS/MS analysis

LC-MS separation was performed on an Agilent ZORBAX RRHD Eclipse Plus C18 2.1 x 50 mm, 1.8 µm column mounted on Agilent 1290 Infinity II UHPLC system coupled to a 6546 qToF mass spectrometer. Solvent composition was: mobile phase A (H_2_O, 0.1% formic acid) and B (methanol, 0.1% formic acid), and the column compartment was kept at 45°C. Five microlitres of sample was injected at 5% B and 0.6 ml/min flow, followed by a linear gradient to 95% B over 3.5 min and 0.6 ml/min. The qTOF instrument (Agilent 6546) was operated in positive scanning mode (50–1,000 m/z) with the following source parameters: VCap, 3,500 V; nozzle voltage, 2,000 V; gas temperature, 225 °C; drying gas 13 L/min; nebulizer, 20 psig; sheath gas temperature 225 °C; sheath gas flow 12 L/min; fragmentor, 130 V. Online mass calibration was performed using a second ionization source and a constant flow (1 mL/min) of reference solution (121.0509 and 922.0098 m/z). The retention times of chemicals were identified by measuring pure chemical standards diluted in deionized water using MassHunter Qualitative Analysis Software (Agilent, version 10.0). Peak integration was performed in MassHunter Quantitative Analysis Software (Agilent, version 10.0) based on known retention time and accurate mass measurement of chemical standards with mass tolerance = 20 ppm, peak filter at signal-to-noise ratio = 2, and retention time tolerance of 0.5 min. Peak area tables were exported as ‘.csv’ files and further analyzed in RStudio (version 4.2.0) for plotting and statistical analyses. LC– MS/MS measurements were performed using the chromatographic separation and source parameters described above, and the auto-MS/MS mode of the instrument with a preferred inclusion list for parent ions with 20 ppm tolerance, iso width set to ‘narrow width’ and collision energy to 20eV. Untargeted metabolomics analysis was performed as described before^8^. In brief, the raw datafiles were aligned in Agilent MassHunter Profinder (10.0) with mass tolerance of 0.002 Da or 20 ppm and retention time tolerance of 0.15 min or 2%. The ‘.cef’ files were exported and used in Mass Profiler Professional to search for ions within m/z 50-300. Identified features were filtered by occurrence frequency of 75% (3 out of 4 replicates). Other parameters used were as default in the software. Moderated t-test was used to calculate p-values, which were corrected for multiple hypothesis testing using the Benjamini-Hochberg method.

### Identification of metabolized mutagens

For each bacterial species, compound fold changes were calculated between time point 12 h and 0 hr and between bacteria-containing samples and control (media-only) samples at 0h in the 4 pools that contained a specific mutagen. Statistical significance of the mutagen intensity differences was assessed with two-sided t-test and p*-*values were FDR-corrected for multiple hypotheses testing. To account for fast mutagen metabolism (within seconds after exposure), fold changes to control at time point 0 were used for mutagen and species combinations for which (i) log2(fold change to control at t = 0) < −5; (ii) FDR-corrected p*-*value (fold change to control at t = 0) < 0.05. To account for variability in chemical measurements, for each drug an adaptive fold-change threshold was calculated as either (75%) or mean + 2 s.d. of the fold changes, for which log2(fold change at t = 12 hr to t = 0 hr) > 0, to account for measurement noise, whichever was greater. Hierarchical clustering was performed with the dendrogram function in R using Euclidean distance between the drug fold-change vectors for each bacterial species.

### Functional chemical group analysis

The occurrences of functional groups were obtained from the rdkit.Chem.Fragments module (Open Source Chemoinformatics, http://www.rdkit.org). Functional group enrichment among metabolized chemicals was calculated using a Fisher’s exact test.

### Germ-free mouse experiments

All mouse experiments were approved by the EMBL’s IACUC (permit number 21-002_HD_MZ). Germ-free C57BL/6J mice of 6-8 weeks of age were kept in flexible plastic gnotobiotic isolators (CBC) with a 12-h light/dark cycle and germ-free status was monitored by PCR and culture-based methods. One group of mice was inoculated with *B. uniformis* via oral gavage of 200 uL of an overnight culture grown in MGAM liquid medium under anaerobic conditions (see above). After four days, bacterial loads in the intestine were determined by CFU plating on BHI blood agar before 3-MeO-AAB was orally administered to the mice at 400 uM via drinking water. Mice were euthanized seven days after chemical exposure, luminal content of small intestine, cecum and colon were collected into sample tubes and snap-frozen and stored at −70 °C until further processing and analysis by LC-MS.

## Data availability

Raw data of the LC-MS analysis have been deposited in the Metabolights public repository with the accession number MTBLS2524. All other data are available in the main text or the supplementary materials.

## Acknowledgement

We thank the Zimmermann lab for helpful discussions, the ChemCore for preparing the chemical pools and measurements of some chemical standards, S. Kasper for suggesting analysis and writing the code for functional group-dependent metabolism, and E. Mastrorilli for providing the original scripts for MassHunter data analysis. This work was supported by the Lung Cancer Research Foundation and the European Molecular Biology Laboratory. M.Z. is an ERC investigator.

---

**Supplementary Figure 1.**
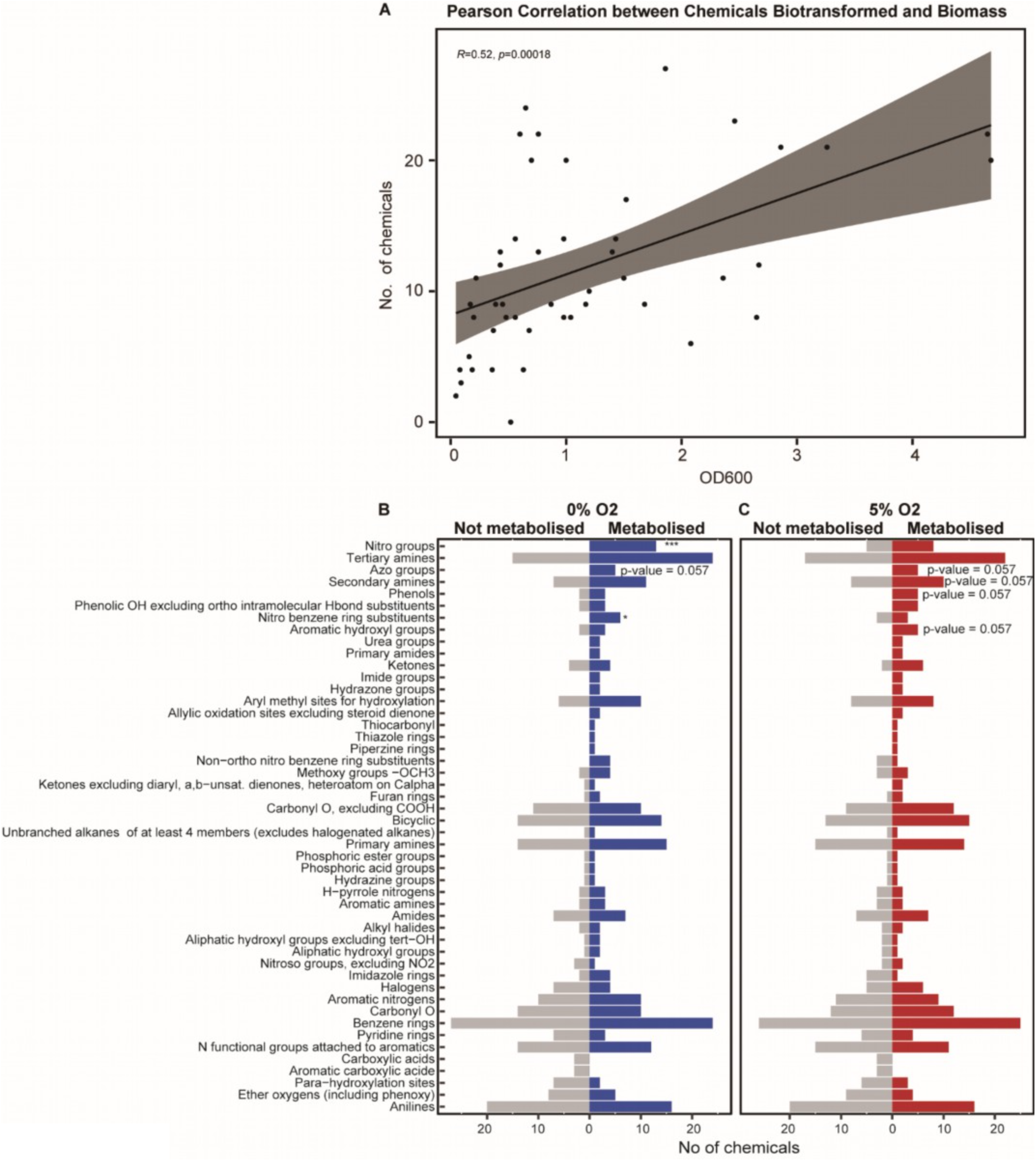
Gut bacterial biotransformation in relation to culture densities and functional groups of carcinogens. **A.** Pearson correlation between total number of chemicals biotransformed by each bacterial species and their biomasses estimated by OD600 measurements. **B-C.** Number of compounds metabolized (**B**: 0% O_2_ = blue, **C**: 5% O_2_ = red) and not metabolized (grey) when carrying different functional groups. Fisher’s exact test was performead from which p-values were obtained for each functional group (Table S4). *, p-value < 0.05; *** p-value < 0.001.

**Supplementary Figure 2.**
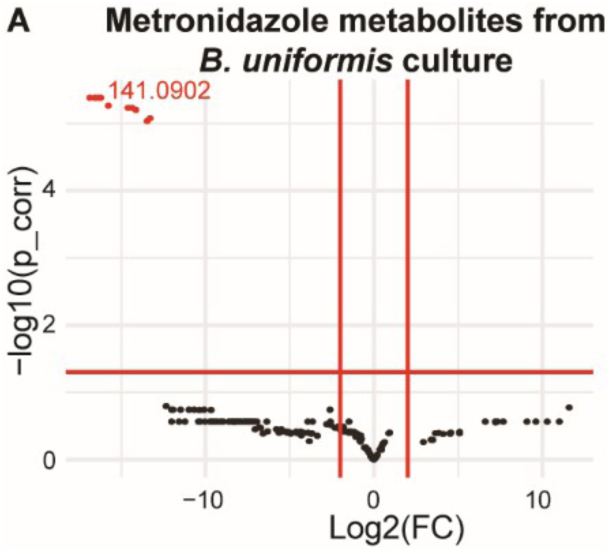
Untargeted metabolomics identified nitro-reduced metabolite of metronidazole by *B. uniformis.* **A.** Volcano plots with p-value (corrected) cutoff 0.05 and Log2(FC) cutoff 2 for untargeted metabolomics analysis of metabolites (neutral masses shown) identified with Agilent MPP software (version 15.1) in metronidazole reaction mixtures after 12 hours of incubation with *B. unifomis* culture. Moderated t-test was used to calculate p-values and Benjamini-Hochberg method was used to correct for multiple hypothesis testing (Tables S15).

**Supplementary Figure 3.**
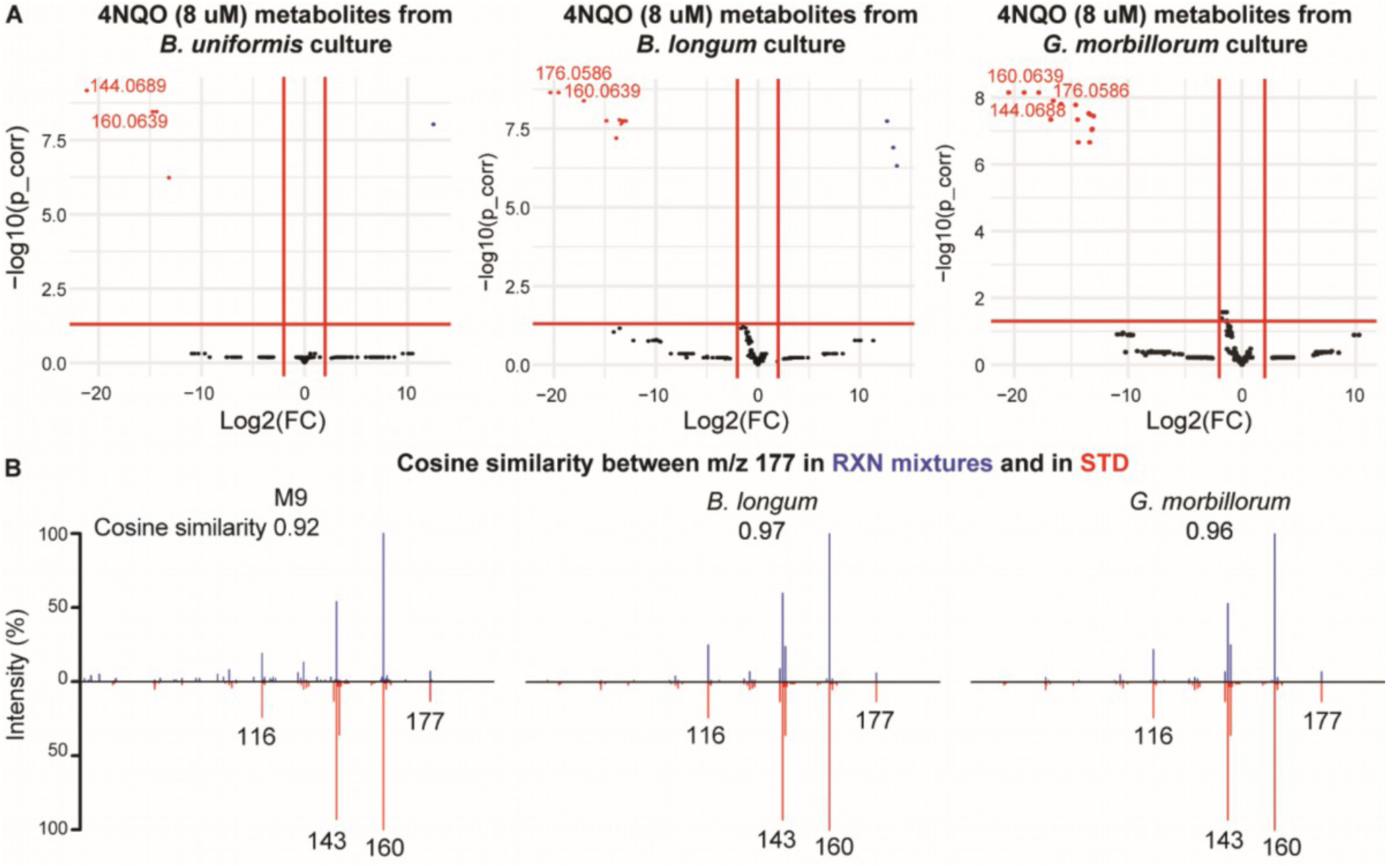
LC-MS/MS spectra of 4-nitroquinoline-1-oxide (4NQO) *N-*hydroxylation metabolite, 4-hydroxyaminoquinoline-1-oxide (4HAQO). **A.** Volcano plots showing FDR-corrected p-value (cutoff 0.05) and Log2(FC) (cutoff 2) for untargeted metabolomics analysis of metabolites (neutral masses shown) identified with Agilent MPP software (version 15.1) in 4NQO reaction mixtures after 12 hours of incubation with *B. unifomis* culture. Moderated t-test was used to calculate p-values and Benjamini-Hochberg method was used to correct for multiple hypothesis testing (Tables S15). **B.** Below x-axis in red: MS/MS fragment ion spectra of 4HAQO chemical standard (TCI Chemicals) with m/z 116, 143, 160 in positive mode. Above x-axis in blue: MS/MS fragment ion spectra of precursor ion with [M+H] m/z 177.066 from reaction mixtures of the respective bacterial species in Fig. 4. Cosine similarities between the fragment ion spectra of precursor with [M+H] m/z 177.066 from each reaction mixture and the chemical standard was calculated respectively.

**Supplementary Figure 4.**
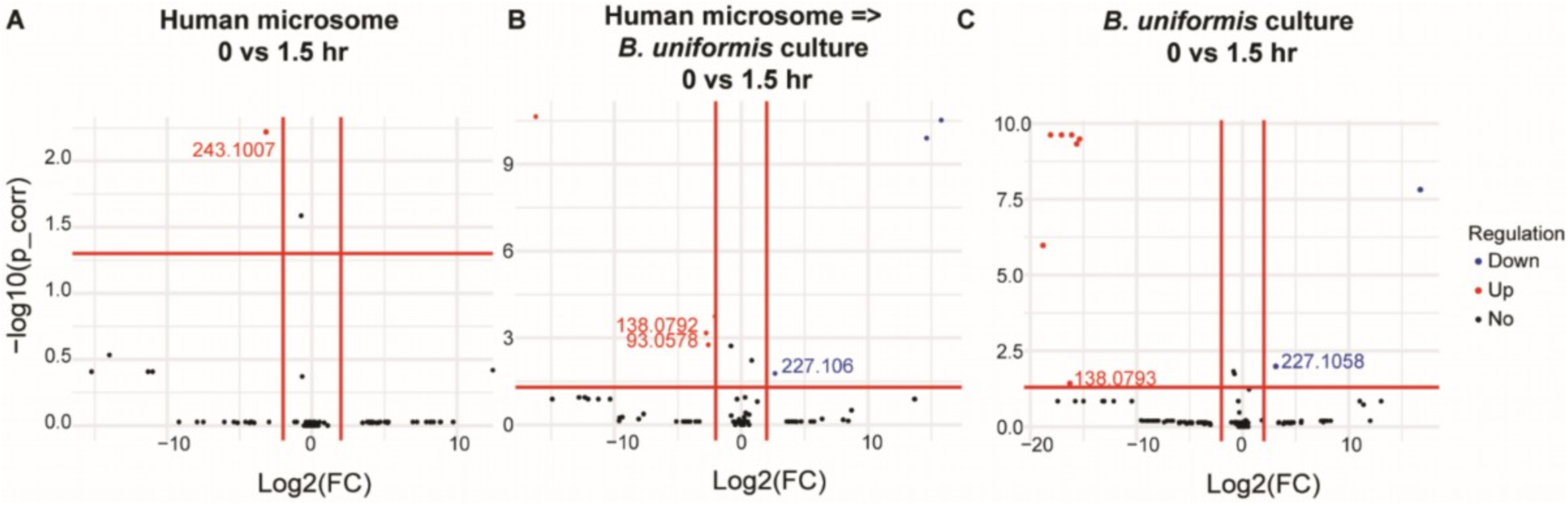
Untargeted metabolomics identifies hydroxylated metabolite of 3-MeO-AAB produced by human microsomes and azo-reduced metabolites by *B. uniformis.* **A-C.** Volcano plots with p-value (corrected) cutoff 0.05 and Log2(fold-change/FC) cutoff 2 for for untargeted metabolomics analysis of metabolites (neutral masses shown) identified with Agilent MPP software (version 15.1) in reaction mixtures before and after 1.5 hr incubation of 3-MeO-AAB with human microsomes (**A**), *B. unifomis* culture (**B**) and *B. unifomis* culture after 1.5 hr incubation with human microsomes (**C**). Moderated t-test was used to calculate p-values and Benjamini-Hochnerg method was used to correct for multiple hypothesis testing (Tables S17-19).

